# Identification of A Potential Inhibitor for Anticancer Target MTHFD2 by Consensus Docking and Molecular Dynamics

**DOI:** 10.1101/2023.11.09.566353

**Authors:** Huiyuan Zhou, Yebin Hong, Floyd A. Beckford

## Abstract

The bifunctional methylenetetrahydrofolate dehydrogenase/cyclohydrolase (MTHFD2) has been recognized as a promising anticancer drug target because it is overexpressed in various types of cancer and is associated with poor prognosis. In the present study, we aimed to discover potential inhibitors from the Enamine HTS library which consists of over one million compounds. A consensus docking-based virtual screening workflow was adopted and two hits, E96 and E41, were identified for being ranked in the top 5 in all docking programs used. To validate the virtual screening result, the binding modes of the two hits were visually inspected with reference to previously published inhibitors B01 and D56, and a similar pattern of binding was observed between the hits and established ligands, indicating the reliability of the docking protocol. The subsequent molecular dynamics simulation and a series of analyses including root mean square deviation, root mean square fluctuation, and radius of gyration reveal that E96 achieved a more stable binding to the receptor than E41. The binding free energy predicted by MM/GBSA calculation confirms E96’s potential to be a potent inhibitor for the target MTHFD2 as it outperforms E41 and the established ligands. In conclusion, this computational study contributes to the drug discovery efforts for the anticancer drug target MTHFD2 by suggesting ligand E96 for further structure-based optimization and *in vitro*/*vivo* experimental validation.

## 1. Introduction

The folate-mediated one-carbon metabolism (FOCM) has been highlighted by previous research to play a crucial role in various biological pathways in organisms that support the proliferation and survival of cells [1, 2]. Moreover, as the rapid growth of cancer cells relies on the synthesis of nucleotide precursors whose carbon units are supplied by the one-carbon metabolism [3], the upregulation of enzymes involved in FOCM has been associated with the emergence of colorectal, lung, breast plus many other types of cancers [4-7]. Among the enzymes involved in FOCM, the bifunctional methylene-tetrahydrofolate dehydrogenase/cyclohydrolase (MTHFD2) which catalyzes the dehydration of 5,10-methylene-THF (CH2-THF) concomitant with the hydrolysis of 5,10-methenyl-THF (CH=THF), stands out as a compelling drug target for its abnormal overexpression in tumor patients with elevated risk of mortality [8, 9]. Recent studies also revealed the non-metabolic role MTHFD2 plays in the upregulation of programmed death-ligand 1 (PD – L1), contributing to cancer immune evasion [10, 11]. As the inhibition of MTHFD2 has exhibited the capacity to curtail the growth of malignant tumors and induce cell death through reduced levels of formate production with minimal side effects [12-14], efforts to discover novel small molecule inhibitors are necessitated.

While traditional drug discovery such as high-throughput screening (HTS) requires exorbitant budgets, computer-aided drug discovery (CADD) has been recognized as an effective strategy to identify druglike compounds among the tremendous chemical space associated with a given target [15]. CADD contributes to drug development by utilizing a suite of computational software that eradicates much of the efforts invested in time-consuming wet lab experiments in early drug discovery stages [16, 17]. Molecular docking and molecular dynamics are arguably the most salient CADD methods widely used in drug discovery nowadays [18, 19]. Most molecular docking programs consist of a search algorithm that varies the position and conformation of a potential drug in a defined pocket on the molecular target which is typically a protein (i.e., a so-called docked pose), and a scoring function that evaluates the binding affinity of sampled docked poses whose scores will be sorted for final ranking [20, 21]. Due to the innate complexity of biochemical reactions *in vivo*, molecular docking cannot be a deterministic approach for recognizing true binders to the target [22].

In contrast, molecular dynamics (MD) attempts to simulate the biological system at an atomic level via physics-based force fields and unveils the behavior of the protein-ligand complex in a solvated environment that complements the oversimplified treatment of the static protein-ligand binding in many molecular docking programs [23]. Furthermore, MD simulations help identify potential drug binding sites and drug binding dynamics and aid in our understanding of intricate interplays in protein-ligand complexes, which would otherwise be challenging to observe with laboratory experiments [24-26].

In this work, a series of *in silico* methods were utilized, including consensus docking, binding mode analysis, MD simulation, and predicted binding free energy by the MM/GBSA method with the goal of identifying potential inhibitors for the anticancer therapeutic target MTHFD2.

## 2. Results

### 2.1 Molecular Docking

Following the 3-stage molecular docking (Figure 1), two compounds (E96 & E41, Figure 2) from the Enamine HTS library were ranked among the top 5 across all the docking programs in view of their binding energy. While E96 (Figure 2A) achieved the first place unanimously among the Vina family, E41 (Figure 2B) was recognized as the best by AutoDock GPU (Table 1) and conversely E96 fell to the fourth place. Due to this inconsistency, both E96 and E41 were retained for further investigation.

**Table 1.** All values are shown as the binding energy (kcal/mol) and ranking (out of 1370 compounds) in each docking program.

| Compound | Enamine ID | AutoDock GPU | AutoDock Vina | Vinardo | QuickVina2 | QuickVinaW |
| --- | --- | --- | --- | --- | --- | --- |
| E96 | Z56823642 | -12.30/5 | -12.96/1 | -10.01/1 | -13.0/1 | -12.9/1 |
| E41 | Z56823893 | -12.56/1 | -12.63/2 | -9.59/2 | -12.6/2 | -12.6/2 |

**Figure 1.**
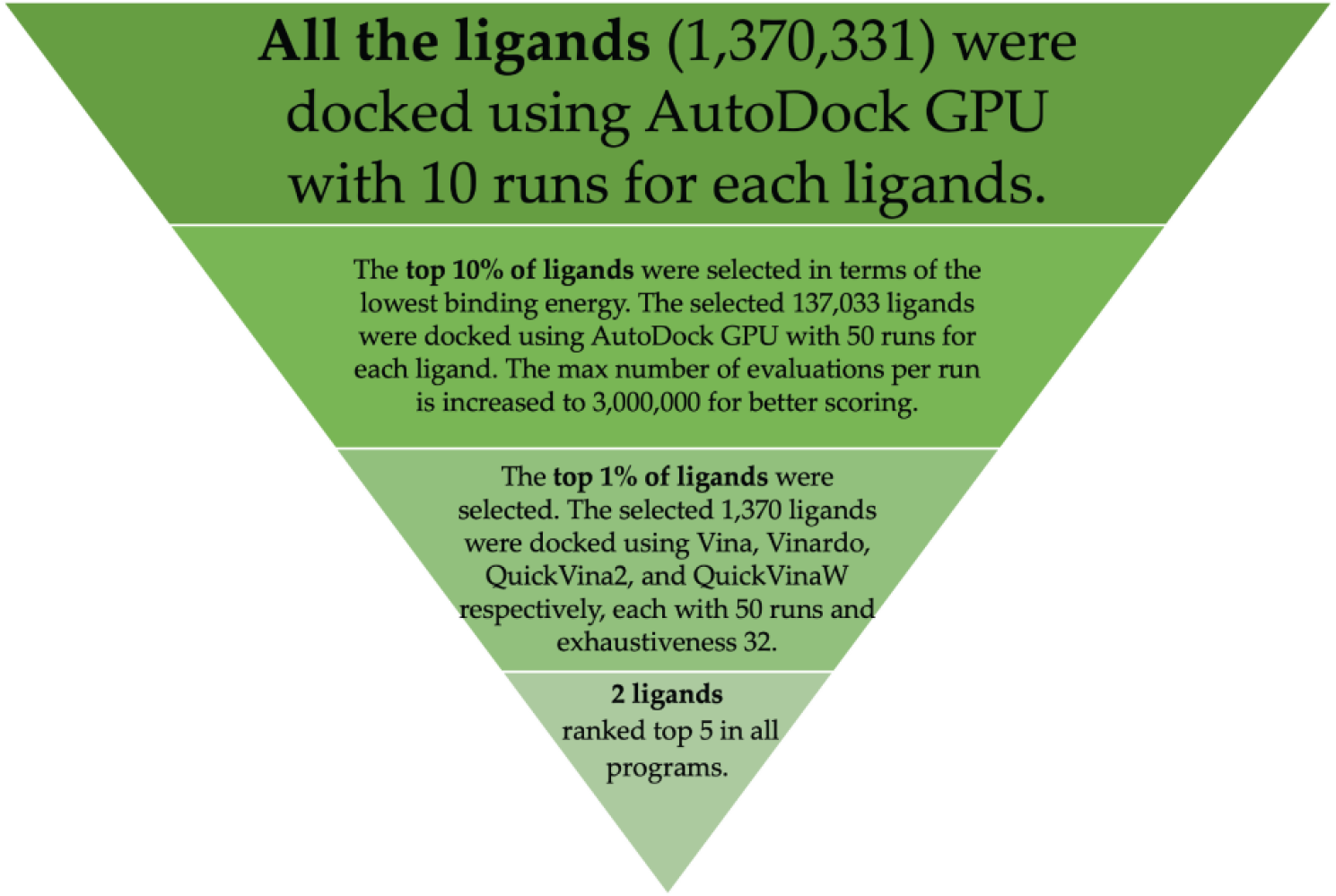
The molecular docking-based virtual screening workflow in this study.

**Figure 2.**
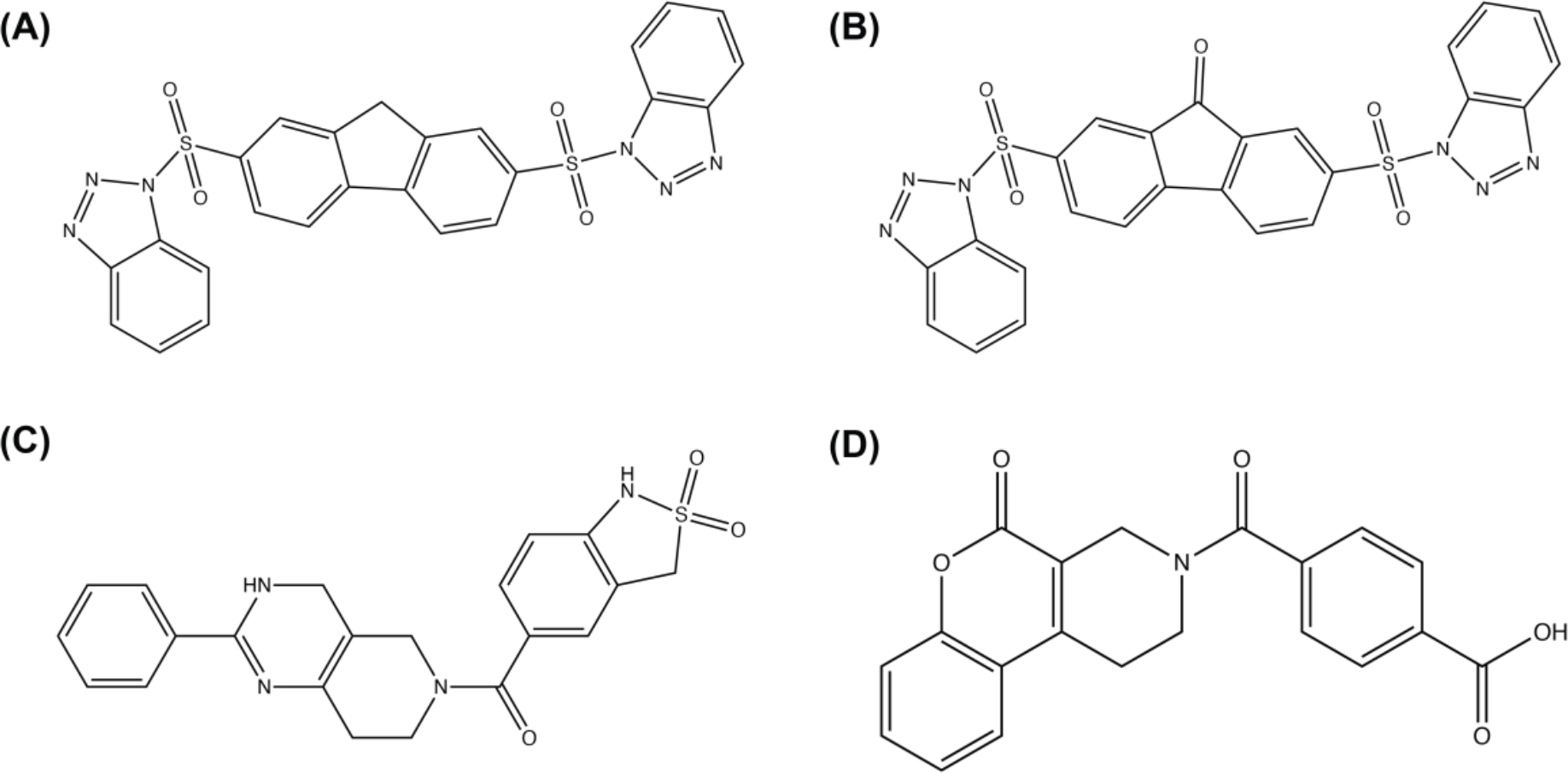
The structures of MTHFD2 inhibitors presented in this study, (A) E96 and (B) E41, and reported substrate site inhibitors of MTHFD2, (C) B01 (compound 1 in ref. 27) and (D) D56 (compound 41 in ref. 27).

Interestingly, while all other programs generated similar binding energies for the two candidates, the Vinardo program generated a significantly different set of numbers.

### 2.2 Binding mode analysis

The binding modes of E96 and E41 were analyzed (Figure 3 and Table 2). Both compounds bind to the folate-binding site of MTHFD2, similar to the previously identified MTHFD2 inhibitors D56 and B01 [27]. Both E96 and E41 formed 3 hydrogen bonds in the binding site of MTHFD2: (1) Lys 88 with an oxygen atom of one sulfonamide group, (2) Gln 132 with the other oxygen atom of that sulfonamide group, and (3) Asn 204 with the backbone nitrogen of the triazole. Tyr 84 and the two aromatic rings of fluorene in both E96 and E41 show a π−π interaction. Another π−π interaction is observed between Phe 157 with the aromatic ring of one benzotriazole in both compounds. For E96, there is an additional π−π interaction with Gly 313 and one aromatic ring of the fluorene. For both ligands, the hydrophobic residues such as Phe 157 and Ala 175 are forming hydrophobic interactions with the aromatic ring of the benzotriazole.

**Table 2.** Key amino acids and their interactions with each compound E96 and E41.

| Compound | Residue in contact | Interaction type |
| --- | --- | --- |
| E96 | Lys 88 | Conventional hydrogen bond |
|  | Gln 132 | Conventional hydrogen bond |
|  | Asn 204 | Conventional hydrogen bond |
|  | Tyr 84 | Pi-pi stacked |
|  | Phe 157 | Pi-pi stacked |
|  | Gly 313 | Pi-pi stacked |
|  | Ala 175 | Pi-Alkyl |
|  | Asn 87 | Carbon hydrogen bond |
|  | Thr 176 | Carbon hydrogen bond |
|  | Pro 314 | Carbon hydrogen bond |
| E41 | Lys 88 | Conventional hydrogen bond |
|  | Gln 132 | Conventional hydrogen bond |
|  | Asn 204 | Conventional hydrogen bond |
|  | Tyr 84 | Pi-pi stacked |
|  | Phe 157 | Pi-pi stacked |
|  | Ala 175 | Pi-Alkyl |
|  | Asn 87 | Carbon hydrogen bond |
|  | Thr 176 | Carbon hydrogen bond |
|  | Pro 314 | Carbon hydrogen bond |

**Figure 3.**
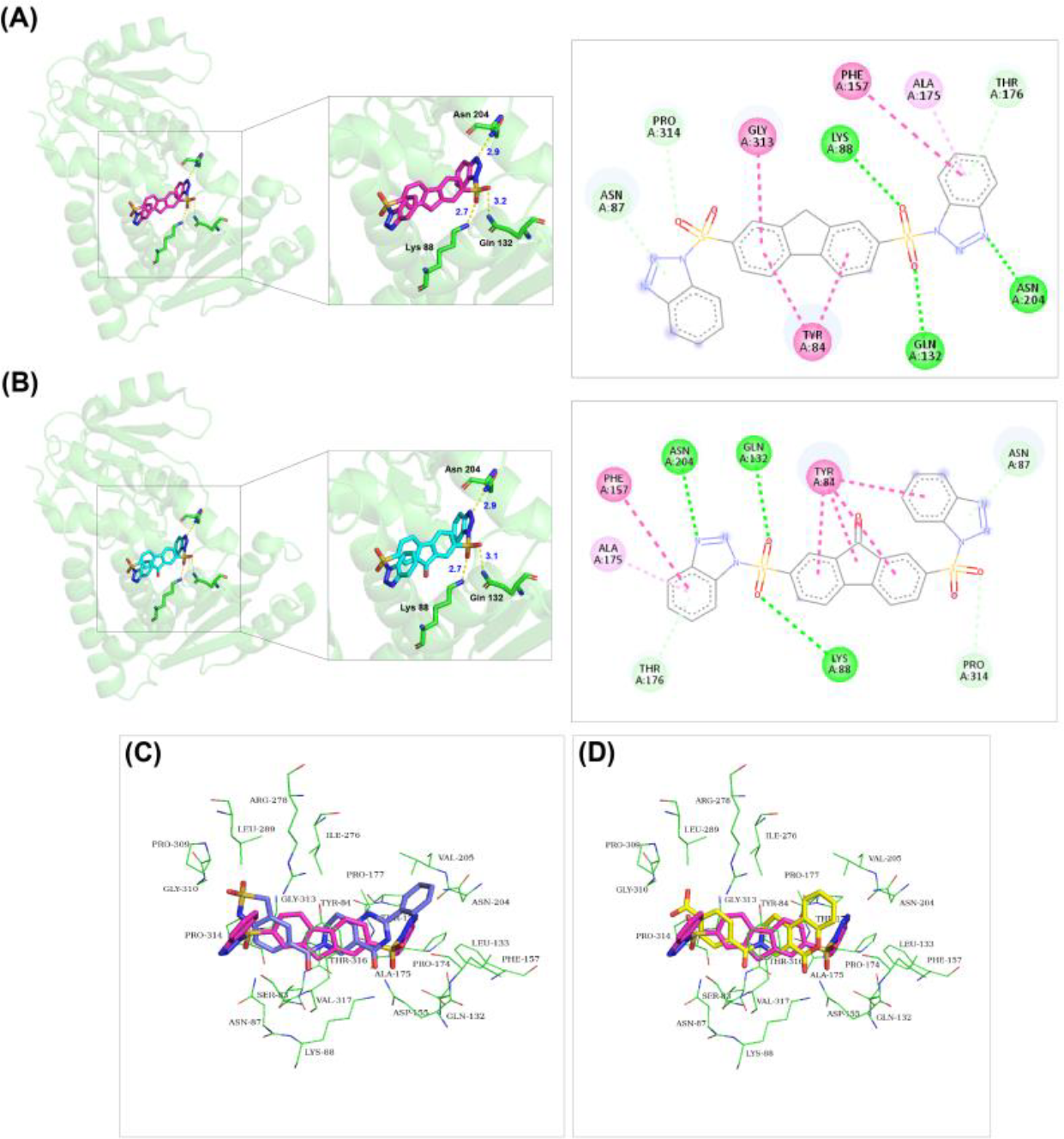
Binding analysis of the (A) MTHFD2-E96 complex and (B) MTHFD2-E41 complex. (C) Superposition of E96 (magenta) and B01 (blue) and (D) superposition of E96 (magenta) and D56 (yellow) with amino acids within 5 Å of the ligand.

### 2.3. Molecular dynamics simulation

The root mean square deviation (RMSD) characterizes the structural deviation in reference to a system’s initial conformation, and a lower RMSD typically indicates a more stable system. From Figure 4A, it can be observed that the protein backbone RMSD of both hits surged to around 0.25 nm soon after the MD simulation was initiated, reflecting adjustments as the protein-ligand complex adapted to the simulation environment. Compared to the protein-E41 complex, the protein-E96 complex exhibited more fluctuation, rising to approximately 0.3 nm at 30 ns and 85 ns. Nevertheless, the protein backbone RMSD of both hits remained similarly consistent, fluctuating from 0.2 nm to 0.3 nm over the course of the simulation, and both converged to around 0.25 nm towards the end. The ligand RMSD of E96 and E41 presents a different trend (Figure 4B). Approximately 10 ns after the simulation was launched, both hits reached a plateau at 0.6 nm. While E96 maintained an equilibrium with subtle variation, the RMSD of E41 started fluctuating at 65 ns, peaking at 1.4 nm around 70 ns. The ligand RMSD of E41 leveled off to 0.6 nm after the fluctuation, but it still oscillated between 0.6 nm and 0.8 nm and displayed greater variability than that of E96 for the remainder of the simulation.

**Figure 4.**
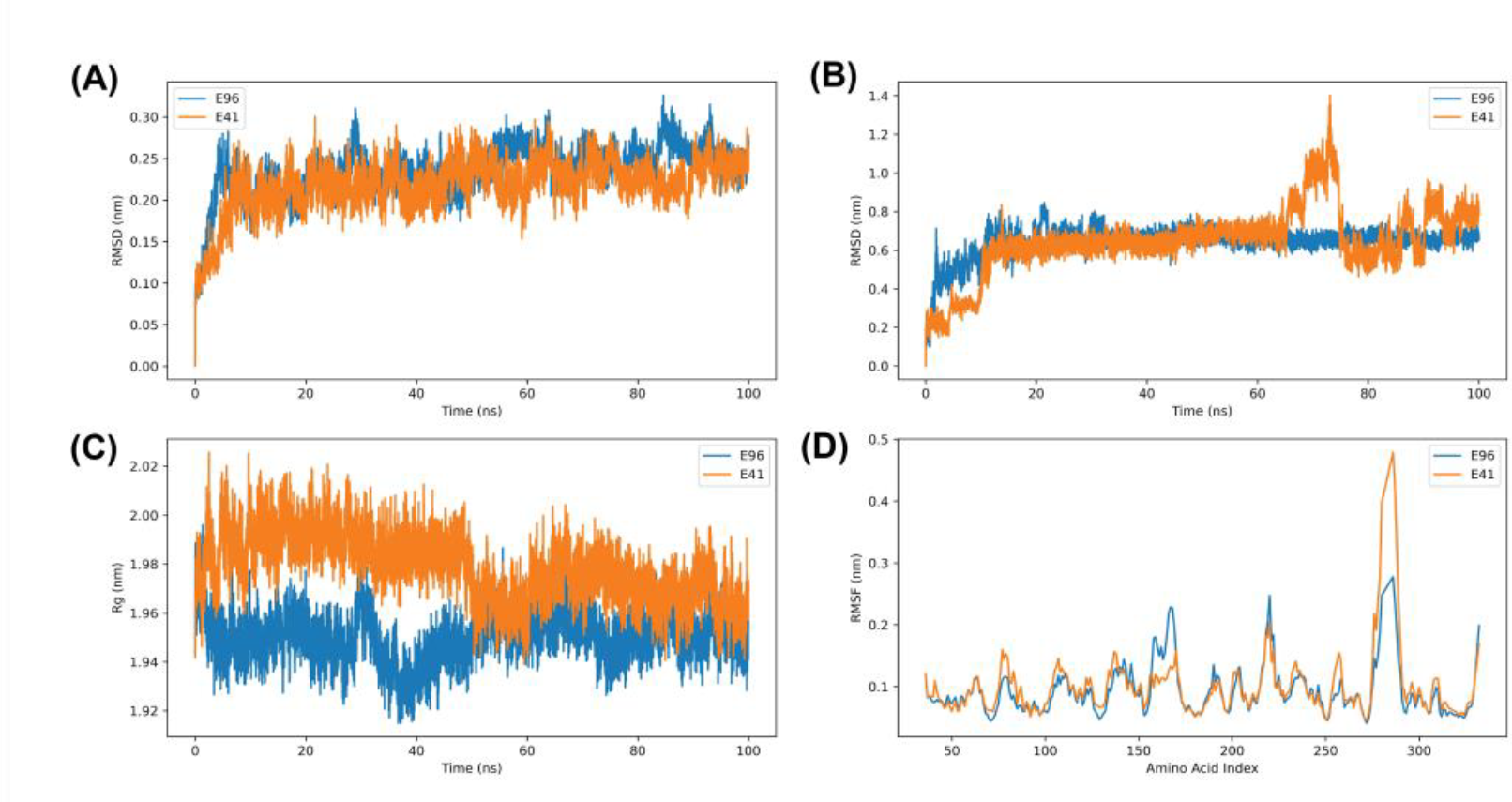
(A) Protein backbone RMSD of E96 and E41. (B) Ligand RMSD of E96 and E41. (C) Radius of gyration (Rg) of E96 and E41 (D) RMSF of amino acid residues of E96 and E41.

The radius of gyration (Rg) reflects the compactness of the protein, and the fluctuation of the Rg value signifies the folding/unfolding of the protein during the simulation [28]. The protein-E41 complex consistently exhibits higher Rg values than the protein-E96 complex, The Rg of the protein-E96 complex (Figure 4C) started decreasing at the beginning of the MD simulation which implies that the protein might have folded in response to the binding of the molecule. By contrast, the Rg trend of the protein-E41 complex diverged from the E96 complex as it rose to ∼2 nm first and remained at a higher value for most of the trajectory. It started declining around the midpoint of the simulation, and ultimately reached a range between 1.96 and 1.98 nm. Both complexes show a significant dip in their Rg values between 40 – 60 ns, indicating a shift towards a more compact structure.

The average mobility of amino acid residues of the protein-ligand complexes was also monitored using root mean square fluctuation (RMSF) as the metric. Both protein-ligand complexes manifest congruent RMSF distribution (Figure 4D), characterized by multiple peaks indicative of regions displaying enhanced flexibility. The E41 complex had a more pronounced fluctuation for residue range 286 – 290 as compared to the E96 complex, which suggests that E41 induced greater mobility within that area. On the contrary, E96 induced more mobility within regions 218 – 221 and 157 – 167. Both complexes show lower RMSF for regions 82 – 101, 180 – 213, 230 – 272, and 298 – 324, depicting residues of more rigidity within these areas.

### 2.4. MM/GBSA calculation

Table 3 presents the binding free energy predicted by MM/GBSA for both two hits and the two established ligands. Between the established ligands, D56 demonstrates a lower binding free energy value (-38.72 ± 1.47 kcal/mol) compared to B01 (-28.57 ± 1.60 kcal/mol), aligning with the order of their experimental IC50 values [27].

**Table 3.**
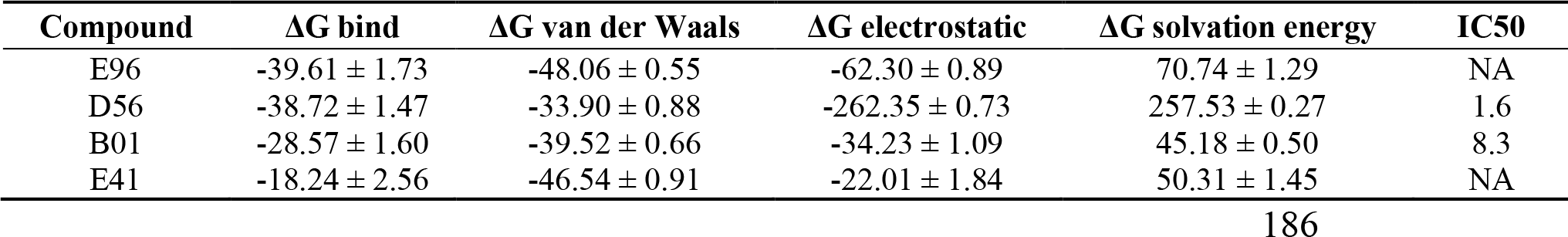
Predicted binding free energy by MM/GBSA method. All energy values are shown in mean value ± standard deviation and are in kcal/mol. IC50 values are in μM.

Furthermore, E96 exhibits a superior binding energy value (-39.61 ± 1.73 kcal/mol) in contrast to E41 (-18.24 ± 2.56 kcal/mol) which is consistent with their performance in the docking stage. E96 also outperforms both established ligands D56 and B01, showing better binding affinity to the receptor. Amino acid residues within 5 Å of the ligand were selected and monitored during the MM/GBSA calculation for the per-residue energy decomposition analysis (Figure 5). For the E96 complex, Arg 278, Ile 276, Lys 88, Tyr 84, and Leu 289 are the top 5 contributing amino acids involved in the ligand binding, highlighting their pivotal role in stabilizing the protein-ligand complex. Conversely, a slightly different set of amino acid residues, namely Tyr 84, Arg 278, Phe 157, Leu289, and Leu 133 exhibited important contributions to the binding of E41. Though Try 84 and Arg 218 are critical in both complexes, they contributed dominantly via electrostatic interaction for the E96 complex but via van der Waals interaction for the E41 complex.

**Figure 5.**
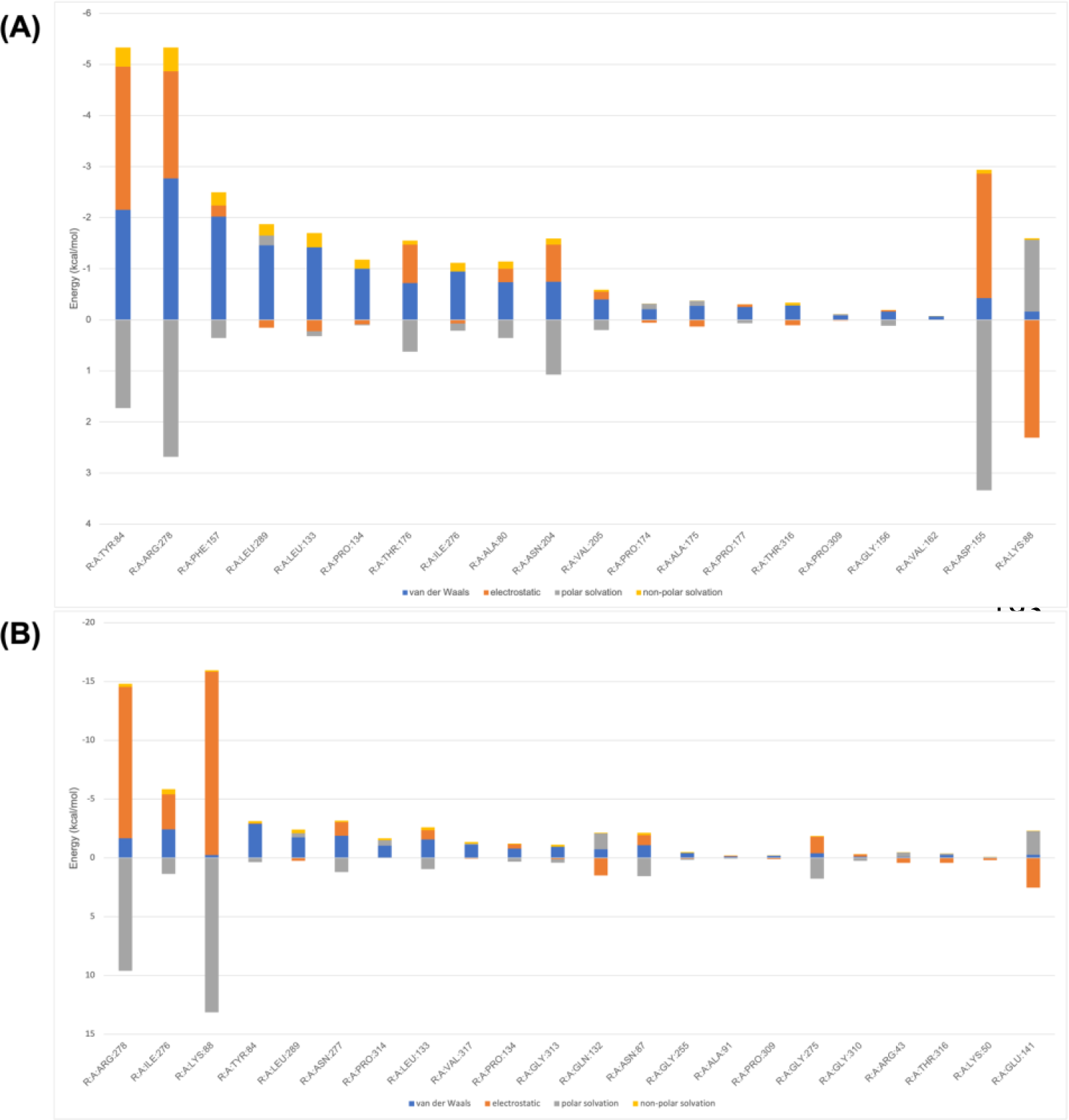
Per-residue energy contribution decomposition of (A) E41 and (B) E96.

## 3. Discussion

As of 2022, it was estimated that more than 10^63^ molecules are drug-like, and over 50 billion molecules could be synthesized upon request, supplied by major chemical vendors [29, 30]. Molecular docking-based virtual screening and advancements in computing power have enabled us to explore vast subsets of the ever-growing chemical space that consist of millions or even billions of molecules for lead compounds, which would otherwise be impossible to realize using traditional biochemical high throughput screening [31]. Despite its successes and prospects, the occurrence of false positives (i.e., ligands that are recommended by docking programs but do not bind experimentally), has impeded drug discovery efforts based on molecular docking [20, 32]. The identification of false positives is sometimes attributed to most docking software’s crudeness in binding affinity prediction and conformation search.

In an attempt to reduce the chance of reporting false positives, the idea of consensus docking was applied in our research which involves five different docking programs [33, 34]. The five docking programs are configured differently in terms of either their search algorithms that explore possible conformations, or their scoring functions that evaluate the binding affinity. While AutoDock GPU adopts the semiempirical free-energy force field-based scoring function [35], the Vina family implements an empirical scoring function [36, 37]. It was then assumed that there should be a certain variation in the final ranking of the 1370 compounds between AutoDock GPU and the Vina family. A post-hoc examination of the docking score ranking corroborates this assumption in that E96 and E41 remain the only two ligands recognized by all programs in the top 5 (top 0.36%, 5/1370) until the search scope is extended to the top 40 (top 2.92%, 40/1370) (Figure S2). This finding adds to our confidence in further analyses of both hits.

As demonstrated in prior research, the analysis of the docked conformations of virtual screened ligands, when compared to experimentally confirmed binding poses, serves as a valuable validation step for assessing the reliability of the docking protocol [38]. The binding mode examination reveals that both ligands occupy the binding pocket of the established ligands B01 and D56, but present slightly different binding modes. Moreover, the hydrogen bonds with Lys 88 and Gln 132 and π−π interaction with Tyr 84 for the two hits were also observed for both the established ligands with MTHFD2, and they are confirmed as essential interactions crucial for the binding [39]. The hydrophobic interactions between the residues (i.e., Phe 157 and Ala 175) and the aromatic ring of the benzotriazole may also play a role in stabilizing the protein-ligand complex.

The MD simulation of the two virtually screened ligands provides further insights into the thermodynamic behavior of the protein-hit complexes in a simulated biological environment. Notably, both protein-hit complexes show acceptable conformational changes during the simulation as evidenced by the average protein backbone RMSD (< 0.3 nm). The juxtaposition of the ligand RMSD trajectories for E96 and E41 underscores their differential behaviors during simulation. E41’s radical fluctuation around 65 ns might have been caused by unfavorable interactions such as the steric effect between the ligand and surrounding amino acid residues, resulting in great orientational changes. After the spike around 65 ns, the ligand RMSD of E41 leveled off but did not reach equilibrium eventually. It is plausible that amino acid residues proximal to the N-terminal of the protein played a critical role in partially stabilizing E41, as indicated by the high RMSF values in the region 286 to 290 in Figure 4D. While conformational flexibility may facilitate ligand binding, excessive mobility can also weaken ligand binding as a result of transient interactions. On the other hand, the highly consistent RMSD of E96 illustrates that the initial docked pose is preferable and well accommodated within the binding pocket throughout the simulation. Regarding the Rg analysis, the generally lower Rg value of E96 showcases that the E96 complex experienced less unfolding and assumed a more compact structure upon ligand binding as compared to E41. The compactness associated with E96 emphasizes the ligand’s potential to foster stronger protein-ligand interactions and reaffirms its stability.

The binding free energy predicted by the MM/GBSA method and comparison with established ligands serve as the ultimate step of validation in this computational study. Remarkably, E96 tops among all examined ligands with predicted ΔG_bind_ = - 39.61 ± 1.73 kcal/mol, suggesting its prospect as a more potent candidate inhibitor compared to the established ligand D56 which has an IC50 equal to 1.6 μM. A closer look at each type of interaction shows that both van der Waals (-48.06 ± 0.55 kcal/mol) and electrostatic (-62.30 ± 0.89 kcal/mol) interactions of E96 contributed substantially and stabilized the complex. By contrast, E41 was mainly stabilized by van der Waals interaction (-46.54 ± 0.91 kcal/mol) and the electrostatic interaction (-22.01 ± 1.84 kcal/mol) contributed far less than that of E96, resulting in its inferior binding affinity. The per-residue energy decomposition both corroborates this result and sheds light on important interacting amino acid residues including Leu 289, Lys 88, and Tyr 84 that are within the top 5 contributing residues for both E96 and E41. Presumably, these residues contributed by van der Waals contact, hydrogen bonding, and π−π stacking interaction respectively as reported previously in Jha *et al*.’s (2023) work [40].

## 4. Materials and Methods

### 4.1. Virtual screening library preparation

The Enamine HTS library was retrieved in 2D sdf format from the Enamine website in March 2023 for its diverse chemotypes [41] (Figure S1). Ten 3D conformations were generated for each ligand using RDKit Python API [42], and further energy-minimized by the built-in MMFF94 forcefield with the lowest energy conformation exported in mol2 format. Ligands in mol2 format were then converted into pdbqt files by the prepare_ligand4.py script offered in the AutoDockTools suite [43].

### 4.2. Molecular docking

Five docking programs were employed in this study: AutoDock GPU, and the Vina family: AutoDock Vina, Vinardo, QuickVina 2, and QuickVina W [16, 35-37, 44-46]. AutoDock GPU was preferred in the first two screening stages for its robust performance due to the utilization of parallel GPU computing [47]. The MTHFD2 structural file was downloaded from the Protein Data Bank (PDB ID: 5tc4) [48] and prepared using AutoDockTools with the addition of hydrogens and the removal of water molecules and ions. For all the docking programs, the grid box was positioned at the center of the crystallized ligand L345899 with the grid size being 20 Å on each side.

In the first stage, the number of runs was set to 10, and the maximum number of asymptotic heuristics evaluations was set to 20,000,000. Ranked by the docking score, the top 10% of ligands were preserved from the initial screening library. For a more refined scoring, the number of runs was increased to 50, and the maximum number of evaluations was increased to 3,000,000 in addition to the parameter adjustment mentioned above in the second screening stage. From this, the top 1% of ligands were picked and then underwent further evaluation by AutoDock GPU and the Vina family. During the consensus docking stage, the number of runs was set to 50 with exhaustiveness being 32 for the Vina family and the parameter setting remained the same for AutoDock GPU.

### 4.3. Binding mode analysis

To visually inspect the binding mode of the hits (i.e., compounds that potentially bind to the target) identified through virtual screening before taking them into further analyses, we selected the docked poses of the two hits predicted by AutoDock Vina as it has been reported in the CASF 2016 benchmark to have the highest success rate of predicting the binding pose in reference to the crystal structure [49]. The two established ligands compound 1 (Figure 2C) and compound 41 (Figure 2D) reported by Kawai *et al*. (2019) [27] were retrieved from PDB ID: 6jid, and ID: 6jib as references. They have been renamed B01 and D56 respectively in our study.

### 4.4. Molecular dynamics simulation

The molecular dynamics (MD) simulations were conducted for the two hits identified through virtual screening and the two reported inhibitors using GROMACS 2023.2 version for 100 ns [50]. For each hit, the docked pose with the lowest predicted binding energy in AutoDock Vina was adopted for the MD simulations, and for the established ligands, their original co-crystallized structures were used. The CHARMM36 forcefield was selected for the MD experiment, and the ligand parameterization files were generated on the CGenFF online server after hydrogens were added using Discovery Studio [51-53]. All systems were solvated in a dodecahedron box with the SPC 216 water model. The appropriate number of Na^+^ or Cl^-^ ions was added to neutralize the solvated system if it was charged. All systems were then energy-minimized with the steepest descent method for a maximum of 50,000 steps. Following energy minimization, all systems were equilibrated under the NVT (isothermal-isovolumetric) and the NPT (isothermal–isobaric) conditions respectively for 100 ps.

### 4.5. Molecular mechanic/generalized-Born surface area (MM/GBSA)

The molecular mechanics with generalized Born and surface area solvation (MM/GBSA) has been a computationally efficient method with decent accuracy to estimate the binding free energy of the protein-ligand complex by breaking down the calculation into the gas phase and the solvent phase [54, 55]:

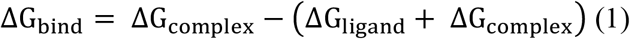

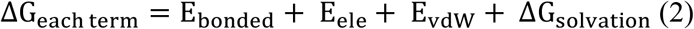

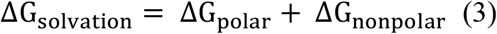

Based on the MD results produced by GROMACS, the 50 - 100 ns trajectory of each tested ligand was sampled with an interval of 10 ps, leading to 5000 frames in total for the MM/GBSA calculation with gmx_MMPBSA 1.6.1, an open-source and free package based on AmberTools [56]. The modified generalized Born model 2 developed by Onufriev *et al*. (2004) was chosen with the addition of 0.15 M salt solution [57].

## 5. Conclusion

In this study, we aimed to discover potential inhibitors targeting the metabolic protein MTHFD2, through molecular docking-based virtual screening and molecular dynamics. Two structurally similar compounds (E96 & E41) were identified from the Enamine HTS library by an ensemble of docking programs. The binding mode analysis demonstrates that both hits form interactions with key amino acid residues in the protein as identified in previous studies. The subsequent molecular dynamics simulation reveals that E96 remains relatively stable over the simulation trajectory and molecular mechanics/generalized Born and surface area (MM/GBSA) calculation affirms this as E96 has a stronger binding affinity with the receptor MTHFD2 compared to the other generated hit E41 as well as the established ligands B01 and D56. These collective findings underscore E96’s prospects for follow-up investigations through structure-based refinement and *in vitro/vivo* validation.

## Author Contributions

Conceptualization, FAB and HZ; data curation, HZ; software, HZ; investigation, HZ, and YH; data visualization, HZ, and YH; writing - original draft, FAB, HZ, and YH; writing – review & editing, FAB, HZ, and YH; All authors have read and agreed to the published version of the manuscript.

## Data Availability Statement

Data in this work can be requested via communication with the corresponding author.

## Acknowledgments

We thank Dr. Mark Spaller from the Division of Natural and Applied Sciences at Duke Kunshan University for his valuable advice on the methodology of this study. We also would like to thank Jasmine Santos and Ethan Mills, senior students at Duke Kunshan University for useful discussions during the project.

