## Supplementary Materials for "Identification of A Potential Inhibitor for Anticancer Target MTHFD2 by Consensus Docking and Molecular Dynamics"

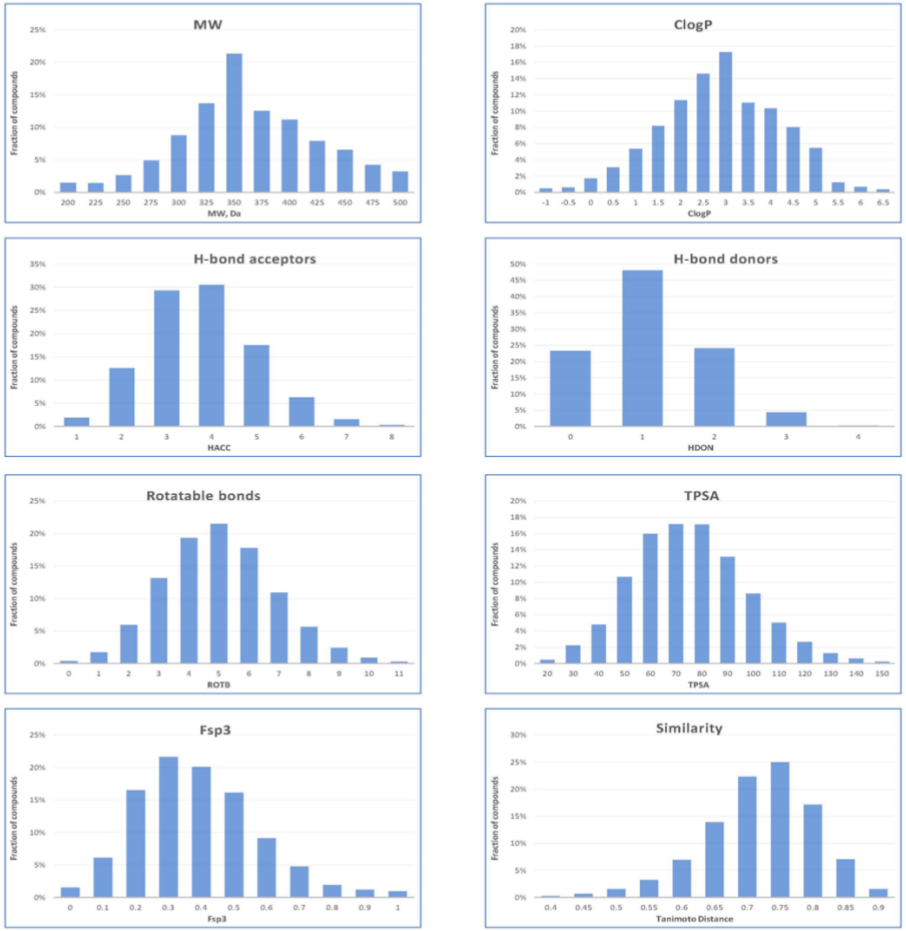


**Figure S1**. The profile of Enamine HTS library. MW: molecular weight; ClogP: evaluation of the partition coefficient; TPSA: topological polar surface area; Fsp3: defined as the number of $sp^3$ hybridized carbon divided by the total number of carbon atoms; The profile was downloaded from https://enamine.net/compound-collections/screening-collection/hts-collection in March 2023.

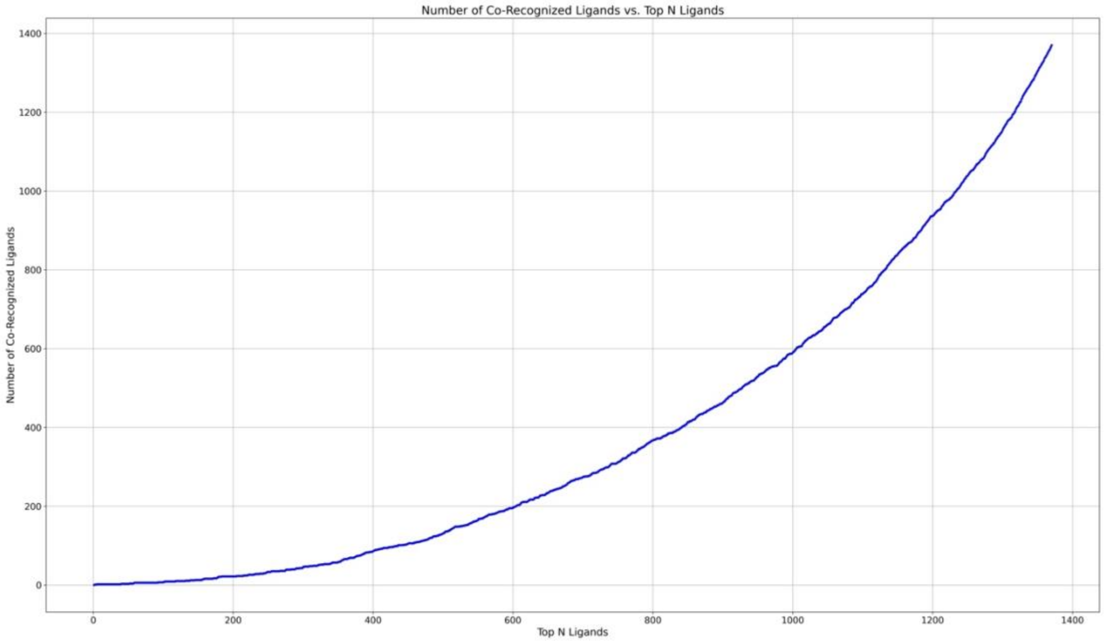


**Figure S2**. The number of co-recognized ligands vs. top N ligands examined.
